# Metabolomic Profiling and Mechanotransduction of Single Chondrocytes Encapsulated in Alginate Microgels

**DOI:** 10.1101/2020.09.28.317008

**Authors:** Jacob P. Fredrikson, Priyanka Brahmachary, Ebru Erdoğan, Zach Archambault, Ronald K. June, Connie B. Chang

**Author notes:** denotes co-corresponding authors.

## Abstract

Human articular cartilage is comprised of two main components, the extracellular matrix (ECM) and the pericellular matrix (PCM). The PCM helps to protect chondrocytes in the cartilage from mechanical loads, but in patients with osteoarthritis, the PCM is weakened resulting in increased chondrocyte stress. As chondrocytes are responsible for cartilage synthesis and maintenance, it is important to understand how mechanical loads affect cellular responses of chondrocytes. Many studies have examined the chondrocyte response to *in vitro* mechanical loading by embedding in stiff agarose. However, these experiments are mostly performed in the absence of PCM which may obscure important responses to mechanotransduction. Here, we demonstrate that drop-based microfluidics allows culture of single chondrocytes in alginate microgels for cell-directed PCM synthesis that closely mimics the *in vivo* microenvironment. Chondrocytes form PCM over 10 days in these single cell microenvironments. Single cell microgels and monolayer controls were encapsulated in high stiffness agarose to mimic the cartilage PCM. After physiological dynamic compression in a custom-built bioreactor, microgels exhibited distinct metabolomic profiles from both uncompressed and monolayer controls. These results demonstrate the potential of single cell encapsulation in alginate microgels to advance cartilage tissue engineering and basic chondrocyte mechanobiology.

## Introduction

Osteoarthritis (OA) is the most common degenerative joint disease which affects over 250 million people worldwide [1]. In osteoarthritis, the articular cartilage, or soft load-bearing tissue that lines the interfaces of joints, begins to break down [2]. This is associated with pain and loss of joint function. The tissue in human articular cartilage is composed of two main components: the extracellular matrix (ECM) and the pericellular matrix (PCM). The cartilage is primarily comprised of ECM, a hydrated matrix of collagen and proteoglycans [2]. The rest of the cartilage consists of PCM which closely encapsulates the chondrocytes cells within the cartilage, like a cocoon, and directly applies stimuli for chondrocyte mechanotransduction [3]. In normal circumstances, cartilage experiences mechanical loads that vary in frequency and amplitude from activities such as walking, running, and jumping [4]. The PCM helps to protect the chondrocytes from these mechanical loads, but in patients with OA, the PCM is weakened resulting in increased chondrocyte stress. [5]. As the chondrocytes are responsible for cartilage synthesis and maintenance, it is important to understand how mechanical loads affect cellular responses of chondrocytes, including their metabolic responses [6].

Many studies have examined the *in vitro* cellular response of chondrocytes to stress by embedding cells in a stiff agarose gel and applying a load to the gel [7-10]. However, these experiments are mostly performed in the absence of PCM which may obscure important metabolic responses involved in mechanotransduction [3]. Chondrocytes *in vivo* are naturally surrounded by a 3-D matrix which has been previously reported to have elastic moduli between 25 and 200 KPa [5, 11, 12]. Three-dimensional agarose models have achieved similar elastic moduli in the range of 20 to 50 kPa with concentrations of 3-5% w/w of low gelling temperature agarose in phosphate buffered saline [9]. However, in these models the cells lack a surrounding PCM and may not accurately represent the *in vivo* functions of the PCM. Thus, the cells do not experience physiologically relevant microenvironments provided by the PCM [3, 9].

Studies have demonstrated that when cultured in millimeter-scale alginate beads, chondrocytes will exhibit phenotypes more similar to *in vivo* conditions [13-15]. These phenotypes include increased glycosaminoglycan (GAG) and collagen synthesis, both of which are components of both the PCM and ECM [14]. In alginate, PCM forms around the cells, yet the elastic moduli of alginate, which is ~ 1 kPA, cannot reach that of previous agarose models which more accurately depict *in vivo* mechanical loads [16-19]. One method of retaining the PCM around the cells is to first grow the cells in a 3-D matrix of alginate and then use various digestive processes to isolate the cells [14]. However, this second digestion may have adverse effects on chondrocytes. Therefore, there remains a need for a technique in which cells can undergo PCM formation in alginate without having a secondary digestion prior to construction of a bulk 3-D matrix. With such a technique, cells can be embedded into a stiffer agarose matrix while retaining properties that better mimic the *in vivo* microenvironment.

One such method is the use of drop-based microfluidics to encapsulate single cells in microscale hydrogels. This approach utilizes fluid flow through small microchannels to create monodisperse picoliter-sized microgels. By flow-focusing an inner alginate hydrogel precursor with an outer oil phase, alginate microgels can be fabricated at kHz rates [19]. This method has been used to encapsulate numerous mammalian cell types [19, 20]. These microgels provide nearly homogenous, highly tunable, 3-D growth environments for cells. Compared to bulk hydrogels, microgels have fewer nutrient limitations and favorable mass transfer characteristics due to the high surface area to volume ratio and small length scale [21]. The individual microgels are easily manipulated through pipetting and centrifugation, allowing for the facile transfer of encapsulated cells between growth environments.

Here, we demonstrate that drop-based microfluidics enables the culturing of single chondrocytes in alginate microgels for cell-directed PCM synthesis that closely mimics the *in vivo* microenvironment. Chondrocytes form PCM over 10 days in these single-cell microenvironments in comparison to cells grown in a monolayer environment which do not form PCM. We investigated the effects of cyclical dynamic compression on cells cultured in microgels and as monolayers. Single-cell microgels and monolayer controls were encapsulated in 4.5 wt% high-stiffness agarose to mimic the cartilage PCM. After physiological dynamic compression in a custom-built bioreactor, we discovered that microgels exhibited distinct metabolomic profiles from both uncompressed and monolayer controls using High Performance Liquid Chromatography – Mass Spec (HPLC-MS) [8]. These microgel-encapsulated chondrocytes develop substantial PCM and have distinct metabolomic profiles compared to monolayer controls. These results demonstrate the potential of single-cell encapsulation in alginate microgels to advance cartilage tissue engineering and basic chondrocyte mechanobiology.

## Methods

### Chondrocyte Harvest and Culture

Primary human chondrocytes were obtained from Stage IV osteoarthritis patients undergoing total joint replacement under IRB approval using established methods [22, 23]. Chondrocytes were isolated by overnight digestion with 2 mg/mL Collagenase Type I (Gibco) for 14 h at 37 °C. Isolated chondrocytes were cultured for 10 days in Dulbecco’s Modified Eagle’s medium (DMEM) (Gibco) supplemented with 10% Fetal Bovine Serum (FBS) (Bio-Techne) and 10,000 I.U./mL penicillin and 10,000 μg/mL streptomycin (Sigma) (hereafter referred to as complete media) in 5% CO_2_ at 37 °C. Chondrocytes were passaged at 90% confluency and seeded at a density of 1 x 10^5^ cells onto 25 x 25 mm microscope coverslips (Fisherbrand). Coverslips with the attached cells were placed in a 60 x 15 mm tissue culture dish containing complete media supplemented with 50 μg/mL L-sodium ascorbate (Sigma) and incubated for 10 days. Media was exchanged every other day. Monolayers grown in complete media lacking L-sodium ascorbate for 10 days at 37 °C were used as controls. Cells were imaged on days 0, 5 and 10 for collagen production. Similarly, SW1353 chondrocytes were seeded on microscope coverslips at a density of 1 x 10^5^ cells and cultured in complete media supplemented with 50 μg/ml L-sodium ascorbate for 10 days in 5% CO_2_ at 37 °C and imaged on days 0, 5 and10. Monolayers grown in complete media lacking L-sodium ascorbate for 10 days at 37 °C were used as controls. SW1353 cells were used between passages 5-20 and primary chondrocytes were used between passages 2-5 post-harvest.

### PDMS Microfluidic device fabrication

Polydimethylsiloxane (PDMS) microfluidic devices were fabricated using standard soft-lithography techniques [24]. Negative master molds were fabricated on 3-in diameter silicon wafers (University Wafer ID: 447) using UV crosslinked Nano SU-8-100 photoresist (Microchem) patterned with photomasks printed on high-resolution transparent plastic film (CAD/Art Inc.). Two-component Sylgard 184 PDMS (Dow SYLGARD 184) was mixed at a 10:1 ratio by mass, poured over wafers in a petri dish (Fisherbrand), and degassed. The devices baked at 55 °C for at least 4 h. The cured devices were cut from the master and ports were punched using a 0.75 mm ID biopsy punch (Well Tech). The devices were plasma bonded to 3 x 2 in glass slides (VWR) for drop makers and 25 x 25 mm type 0 coverslips (VWR) for imaging arrays by exposing the PDMS and glass to oxygen plasma for 1 min at high power, 45 watts, and 700 mTorr oxygen pressure using a plasma cleaner (Harrick Scientific Corp PDC-002). The bonded devices were baked at 55 °C for at least 1 h to increase the strength of the plasma bond. Following baking, drop makers were filled with a hydrophobic treatment (PPG Industries 24 Aquapel) for 5 min, flushed with air, and baked at 55 °C again for 1 h.

### Preparation of Precursor Solutions

*Oil Preparation*: The surfactant Krytox 157 FSH (Miller-Stephenson) was added to the fluorinated oil, HFE 7500 (3M Novec), at 4% w/w. The solution was filtered through a 0.2 μm hydrophobic polytetrafluoroethylene (PTFE) syringe filter (GVS). *Alginate Precursor Preparation (A)*: 80 mM CaCl2 (ACROS Organics), 80 mM Na2EDTA (Fisher Chemical), and 40 mM MOPS (VWR) were mixed with ultra-pure water (18.2 MΩ•cm, MiliQ) to make 50 mL of solution and the pH was adjusted to 7.2 using 2 M NaOH (VWR). Sodium alginate (Sigma, 9005-38-3) was added to 10 mL of the solution at 1.5% w/w and the solution was filtered through a 0.2 μm hydrophilic polyethersulfone (PES) syringe filter (MilliporeSigma Millex). *Alginate Precursor Preparation (B)*: 80 mM Zn(CH_3_CO_2_)_2_ (Alfa Aesar), 80 mM EDDA (TCI Chemicals), and 40 mM MOPS were mixed with ultra-pure water (18.2 MΩ•cm, MiliQ) to make 50 mL of solution and the pH was adjusted to 7.2 using 2 M NaOH. Sodium alginate was added to 10 mL of the solution at 1.5% w/w and the solution was filtered through a 0.2 μm PES syringe filter.

### Cell Encapsulation in Microgels

To prepare cells for encapsulation, cells were removed from monolayers using 2 mL of Trypsin-EDTA and suspended in media. Cells were centrifuged at 500 x g for 5 min (Thermo Scientific ST-16) and washed with 10 mL of PBS twice. In order to minimize the number of empty drops, cells were then suspended in alginate precursor solution ‘*A*’ at 5 x 10^6^ cells/mL (final concentration 2.5 x 10^6^ cells/mL) and loaded into 1 or 3 mL syringes (BD luer lock). Following previous protocols, cells were encapsulated in 100 μm diameter alginate microgels [19]. The continuous flow rate used was *Q*_Oil_ = 2000 μL/h and the dispersed phase flow rates used were *Q*_AlgA_ = *Q*_AlgB_ = 250 μL/h. Microgels were produced in parallel by running two drop making devices for 4 h with the goal of encapsulating at least 1 x 10^7^ cells. After encapsulation and gelation, microgels were washed with 500 μL HFE 7500 and 100 μL 20% v/v 1H,1H,2H,2H-Perflourooctonal (PFO) (VWR) in HFE for each 125 μL of microgels multiple times to remove the surfactant. Microgels were collected from the oil phase with 500 μL PBS and suspended in the appropriate media (see following section).

### Cell Culture in Microgels

Approximately 200 μL of microgels containing ~1.3 cells/microgel were placed in each well plate of a six well plate (Falcon) and covered with 2 mL of media. Cells were either cultured with complete media or complete media supplemented with 50 μg/mL L-sodium ascorbate to promote collagen synthesis. Every 3 days, media was exchanged by collecting the contents of each well, centrifuging at 500 x g for 2 min, removing the supernatant, and suspending the microgels in 2 mL of new media.

### Cell Viability Assay

After 10 days of culture, 100 μL of microgel encapsulated SW1353 cells were mixed with 800 μL of PBS and 100 μL of Trypan Blue (Corning). 10 μL of the solution was pipetted onto a hemocytometer (Reichert). Live and dead cells were counted. Dead cells were stained blue while live cells remain colorless.

### Florescence Imaging and Analysis

SW1353 cells were imaged using an epi-fluorescent microscope (Leica dmi-8) with a 40 X objective over 10 days to capture bright field and auto-fluorescence (ex. 350 nm / em. 470 nm). For autofluorescence imaging, 30 μL of microgels and media were flowed into a microfluidic array each day and 19-23 cells per condition were imaged. The background intensity was subtracted from each image using Fiji Image J and the florescence intensity of each cell was measured using the Fiji ImageJ distribution of TrackMate [25, 26].

### Staining and Confocal Imaging

#### Monolayer Controls

Primary chondrocytes and SW1353 cells grown in monolayers on microscope coverslips were fixed in 4% v/v Paraformaldehyde – 1 X Phosphate Buffered Saline (PBS) for 10 min at room temperature followed by three 5 min washes of 1 X PBS. Cells were permeabilized with a blocking solution containing 0.1% Triton X-100 - 1% Bovine Serum Albumin – 1 X PBS for 30 min at room temperature. Cells were then incubated with the primary antibody to Collagen VI (Rabbit Polyclonal Anti-Collagen VI antibody, ab6588 from Abcam) in 0.1% Triton X-100 - 1% Bovine Serum Albumin - 1 X PBS for 1 h at room temperature. After three 5 min washes with 1 X PBS, cells were incubated with a mixture of the secondary antibody Donkey anti-Rabbit IgG H&L Alexa Fluo^R^488 (Abcam), Vibrant™DyeCycle™ Violet Stain for Nuclei (Invitrogen), and MitoTracker™ Red CMXRos (Invitrogen) for mitochondrial detection in 0.1% Triton X-100 - 1% Bovine Serum Albumin – 1 X PBS for 1 h at room temperature. Cells were washed 3 times with 1 X PBS and the coverslip was mounted on a glass slide with ProLong™ Diamond Antifade Mountant (Invitrogen). Digital images were acquired on a Leica TCS SP8 confocal microscope with the Leica Application Suite Advanced Fluorescence software.

#### Microgel-encapsulated Chondrocytes

Alginate beads with encapsulated chondrocytes were spun down at 500 x g for 2 min, washed with 1 X Phosphate Buffered Saline (PBS), and then fixed with 4% v/v Paraformaldehyde – 1 X PBS for 15 min at room temperature. Paraformaldehyde-fixed microgel-encapsulated cells were permeabilized with a blocking solution containing 1.0% Triton X-100 - 3% Bovine Serum Albumin – 1 X PBS for 30 min at room temperature. Microgels were then incubated with primary antibody to Collagen VI (Rabbit Polyclonal Anti-Collagen VI antibody, ab6588 from Abcam) in 1.0% Triton X-100 - 3% Bovine Serum Albumin – 1 X PBS for 1 h at room temperature. Following three washes of 5 min each with 1 X PBS, encapsulated samples were incubated with a mixture of secondary antibody Donkey anti-Rabbit IgG H&L Alexa Fluo^R^488 (Abcam), Vibrant™DyeCycle™ Violet Stain for Nuclei (Invitrogen), and MitoTracker™ Red CMXRos (Invitrogen) for mitochondrial detection in 1.0% Triton X-100 - 3% Bovine Serum Albumin – 1 X PBS for 1 h at room temperature. Microgel constructs were washed three times with 1 X PBS. 100 μl of the alginate beads were cytospun onto single frosted adhesive slides (Tanner Scientific) using a Thermo Scientific Cytospin™ 4 Cytocentrifuge for immunocytochemistry and mounted with ProLong™ Diamond Antifade Mountant (Invitrogen).

### Encapsulation in High Stiffness Agarose, Mechanical Stimulation, and Metabolite Extraction

After 9 days in culture, microgels and SW1353 cells from each condition were mixed with low melting temperature agarose (Sigma Aldrich) at a final agarose concentration of 4.5% w/w [9, 27]. Final cell concentrations for each condition were 1 x 10^6^ cells/mL with each gel being a 0.5 mL cylinder 12.75 mm in height. The agarose gels were each placed in individual wells of a 12 well plate (Falcon) and cultured with the corresponding media for 2 days. After 2 days, each gel was loaded onto a custom-built bioreactor at 37 °C 5% CO_2_ and 95% relative humidity in PBS and preloaded at 5% strain [8]. After 30 min, 2% cyclic strain at 1.1 Hz was applied for 15 min. Gels were washed with PBS and frozen for 1 min with liquid nitrogen. Gels were then crushed and placed in a −80 °C freezer for 1 h. Gels were removed from the freezer and a 1 mL solution of equal volumes HPLC-grade methanol (Fisher Chemical) and acetone (Sigma Aldrich) was added to each sample. Samples were vortexed every 5 min for 20 min and kept at −20 °C overnight. The next day samples were centrifuged at 13,000 rpm for 10 min at 4 °C (Thermo Scientific Sorvall Legend X1R) The supernatant was removed and placed in a speed-VAC (Savant SC110) for 6.5 h. The resulting pellet was suspended in a 1:1 mixture of HPLC grade water (Fisher Chemical) and acetonitrile (Fisher Chemical) and placed in a −80 °C freezer until used for HPLC-MS.

### Untargeted Metabolic Profiling

Changes in cellular activity were studied using untargeted metabolomic profiling by HPLC-MS (high performance liquid chromatography coupled to mass spectrometry). Chromatography was performed in normal phase with established protocols using a Cogent Diamond Hydride HILIC 150 x 2.1 mm column using an Agilent 1290 UPLC system [28]. Mass spectrometry was performed using an Agilent 6538 Q-TOF. Metabolites with a median intensity of zero across all experimental groups were excluded from analysis.

Undetected remaining metabolites (*i.e*. intensity of zero) were replaced with a value of one half the minimum peak intensity for statistical analyses. Statistical analysis was performed in Metaboanalyst. Data were first log transformed and standardized. Metabolomic profiles between experimental groups were compared using principal components analysis (PCA), hierarchical clustering, volcano plot analysis, and pathway enrichment analysis. Significance was assessed with false discovery rate corrections using a significance level of 0.05 selected *a priori*.

### Rheology

The storage and loss moduli of each gel used was measured using a TA Instruments AR-G2 parallel plate rheometer. Gel mixtures were poured into 2 mL molds and gelled. They were then loaded onto the rheometer and 1% strain was applied from 100 to 0.1 rad/s.

### Statistical Analysis

SW1353 cell viabilities were compared using a Welch Two Sample t-test in R [29, 30]. The rate of autofluorescence increase was calculated using a weighted multiple linear regression in R [29, 30]. Each measurement was weighted by the inverse variance of the mean of each sample.

## Results and Discussion

### Chondrocyte encapsulation in alginate microgels

Previous studies have investigated the effects of alginate macrogels on the formation of pericellular matrix, but while these macrogels support the formation of PCM, they have insufficient stiffness to mimic the *in vivo* elastic modulus surrounding chondrocytes [13-15]. Here, drop-based microfluidics were used to first encapsulate single cells (SW1353 and primary human chondrocytes) in alginate microgels (Fig. 1A). In order to produce alginate microgels in a biocompatible manner, the competitive ligand exchange crosslinking (CLEX) method for crosslinking alginate was used [19]. After 9 days in culture, concentrated microgels were suspended in 4.5% w/w agarose and gelled (Fig. 1B). The agarose constructs were allowed to come to equilibrium in a custom-built bioreactor and then subjected to physiological dynamic compression (Fig. 1C). Following compression metabolites were extracted from the chondrocytes and analyzed using HPLC-MS.

**Figure 1:**
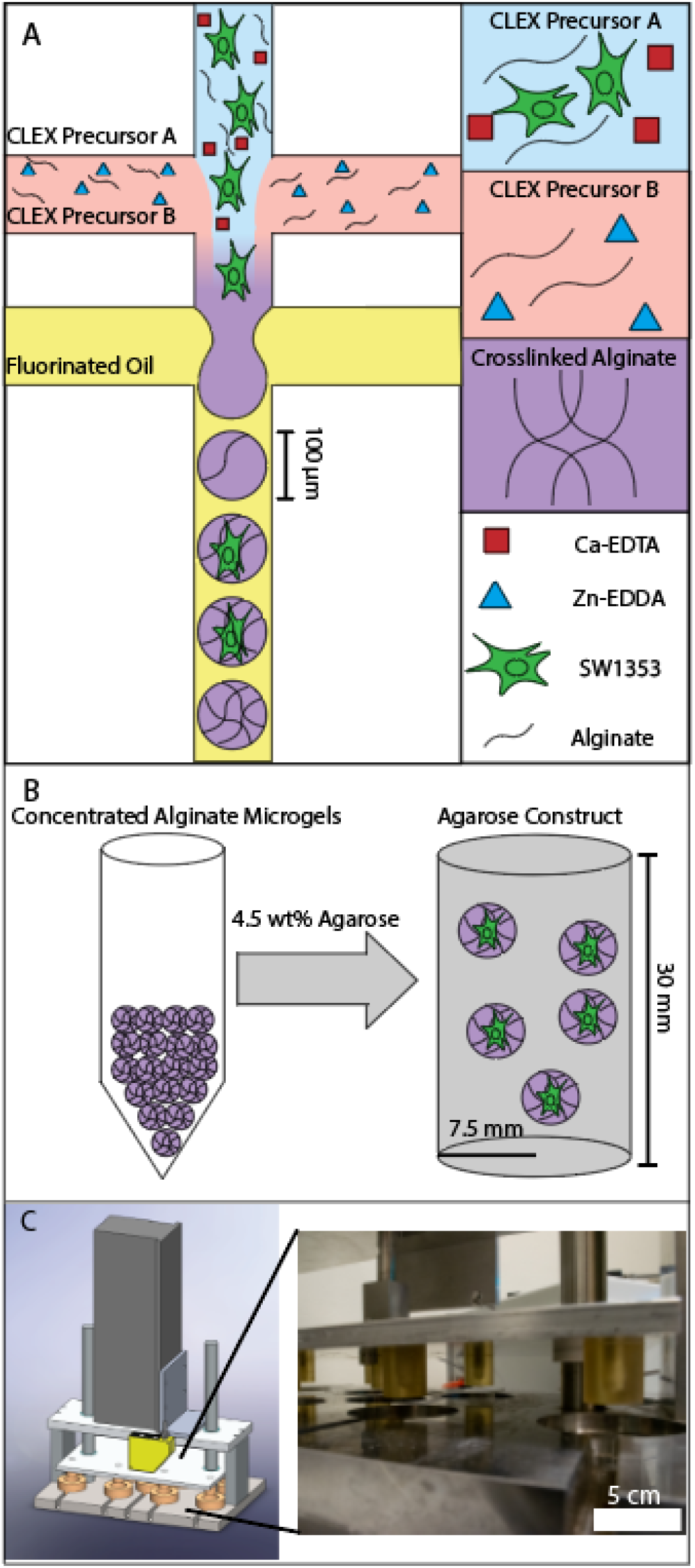
(A) Schematic of alginate microgel production. Cell laden CLEX precursor A is mixed with CLEX precursor B and flow focused with 4% FSH - HFE 7500. (B) After culture for 9 days, concentrated alginate microgels are mixed with agarose to a final agarose concentration of 4.5% w/w and 80% v/v. (C) After 2 days in the agarose, a custom built bioreactor applied 2% cyclic strain after a 5% strain preload. Following 15 min of cyclic compression, metabolites are extracted from the samples and analyzed using HPLC-MS.

### Pericellular matrix formation in alginate microgels

To visualize the PCM formation, SW1353 and primary human chondrocytes were fixed on days 0, 5, and 10, stained for collagen VI and imaged using confocal microscopy. When cultured with ascorbate, the microgel encapsulated primary chondrocytes produced the most collagen VI of all conditions tested (Fig 2). The encapsulated primary chondrocytes produced collagen VI that closely surrounds the outside of the cell as observed using confocal microscopy. By day 5 (Fig. 2 xi), a thin (~1 μm) layer of collagen had formed around most of the cells and by day 10 (Fig. 2 xii) the collagen layer was ~2 to 5 microns thick surrounding the entire perimeter of the cell. When cultured without ascorbate, the encapsulated primary chondrocytes showed similar trends but produced less collagen (SI Fig. 1). The SW1353 cells cultured in microgels showed an increase in collagen VI by day 5, and by day 10, the majority of the cell was covered with collagen VI when ascorbate was present (Fig 2 iv-v). The SW1353 cells however, produced less collagen VI than the primary cells.

**Figure 2:**
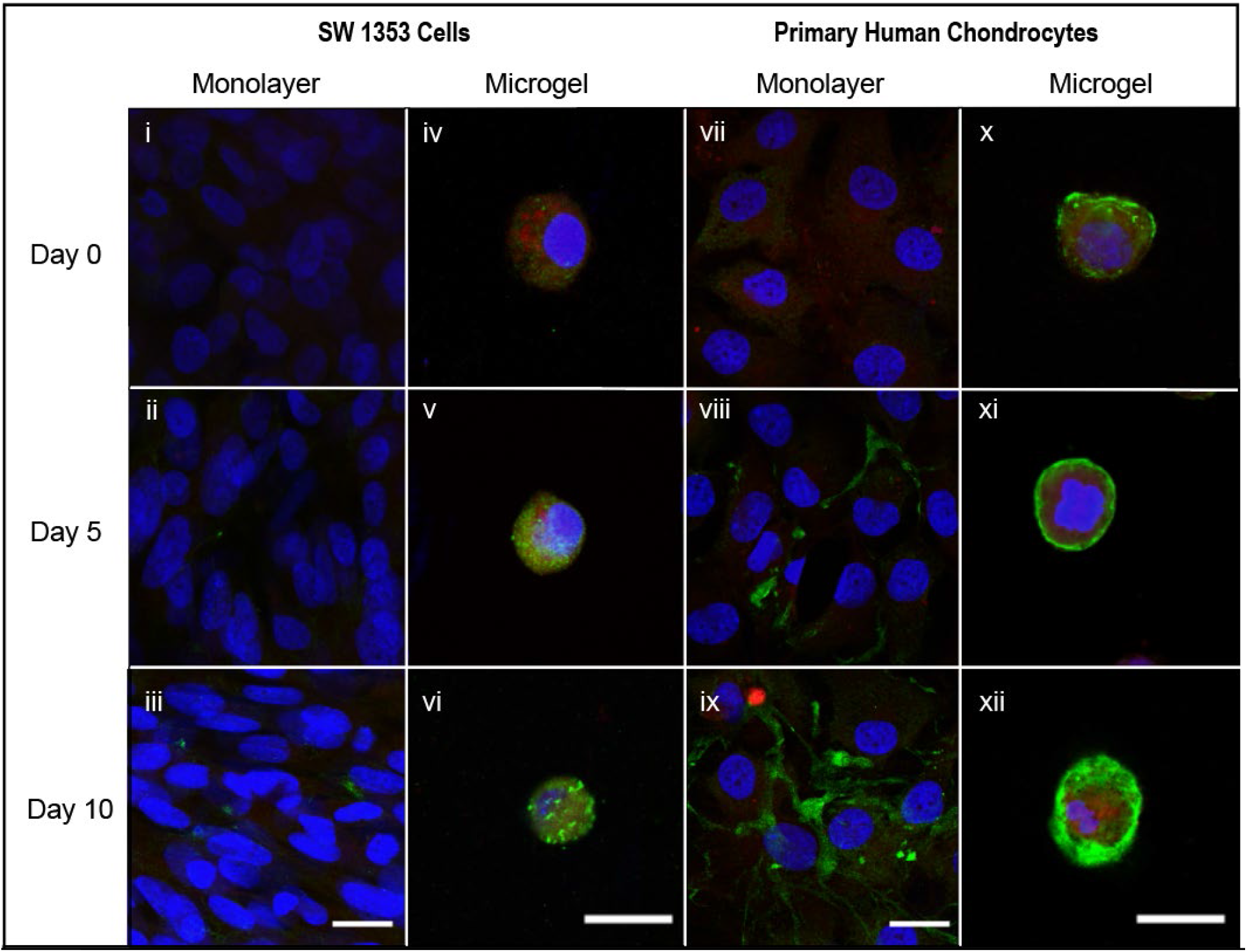
SW 1353 and Human Primary Chondrocyte cells fixed with 4% v/v paraformaldehyde in PBS. Cells were stained for Collagen VI (green), mitochondria (red), and nuclei (blue). Cells were imaged on days 0, 5, and 10 on monolayers and in 1.5% w/w alginate microgels. Cells pictured were cultured in media with 50 μg/mL ascorbate. Scale bar is 20 μm.

SW1353 cells cultured in two-dimensional monolayers showed no collagen VI on day 0 and had negligible collagen VI staining by day 10 when ascorbate was present in the media (Fig. 2 i-iii). When cultured in monolayer without ascorbate, primary chondrocytes produced a small amount of collagen VI whereas the primary chondrocytes cultured with ascorbate produced more (SI Fig. 1 / Fig. 2 vii-ix). While the primary chondrocytes cultured in monolayer produced collagen VI, it was typically located in small concentrated regions and did not surround the cells in contrast to both the microgels and the endogenous PCM.

To quantify collagen formation, cellular auto-fluorescence was quantified using an epi-fluorescent microscope. Collagen, a main component of the PCM, shows substantial auto-fluorescence upon excitation from light ranging from UV to blue [31-33]. Thus collagen-based PCM formation can be quantified over time by repeated imaging of auto-fluorescence in this label-free approach. We validated this method by comparing auto-fluorescence and collagen VI staining in the same samples, and the data showed strong correlations between the two techniques. Each day for 10 days SW1353 cells grown in monolayer and SW1353 cells grown in alginate microgels were imaged (Fig. 3A, SI Fig. 2A, B). The auto-fluorescence signal followed similar trends to the collagen VI staining. The auto-fluorescence signal linearly increased with time in microgels at significantly higher rates than monolayer (p < 2 x 10^-16^, Fig. 3B). For microgel encapsulated chondrocytes, the addition of ascorbate significantly increased the rate at which the auto-fluorescence signal increased (p = 5.20 x 10^-5^). The auto-fluorescence signal closely surrounded the cell similar to the endogenous PCM (Fig. 3A). These data indicate that single-cell encapsulation of chondrocytes within alginate microgels dramatically increases collagen production compared to monolayer controls.

**Figure 3:**
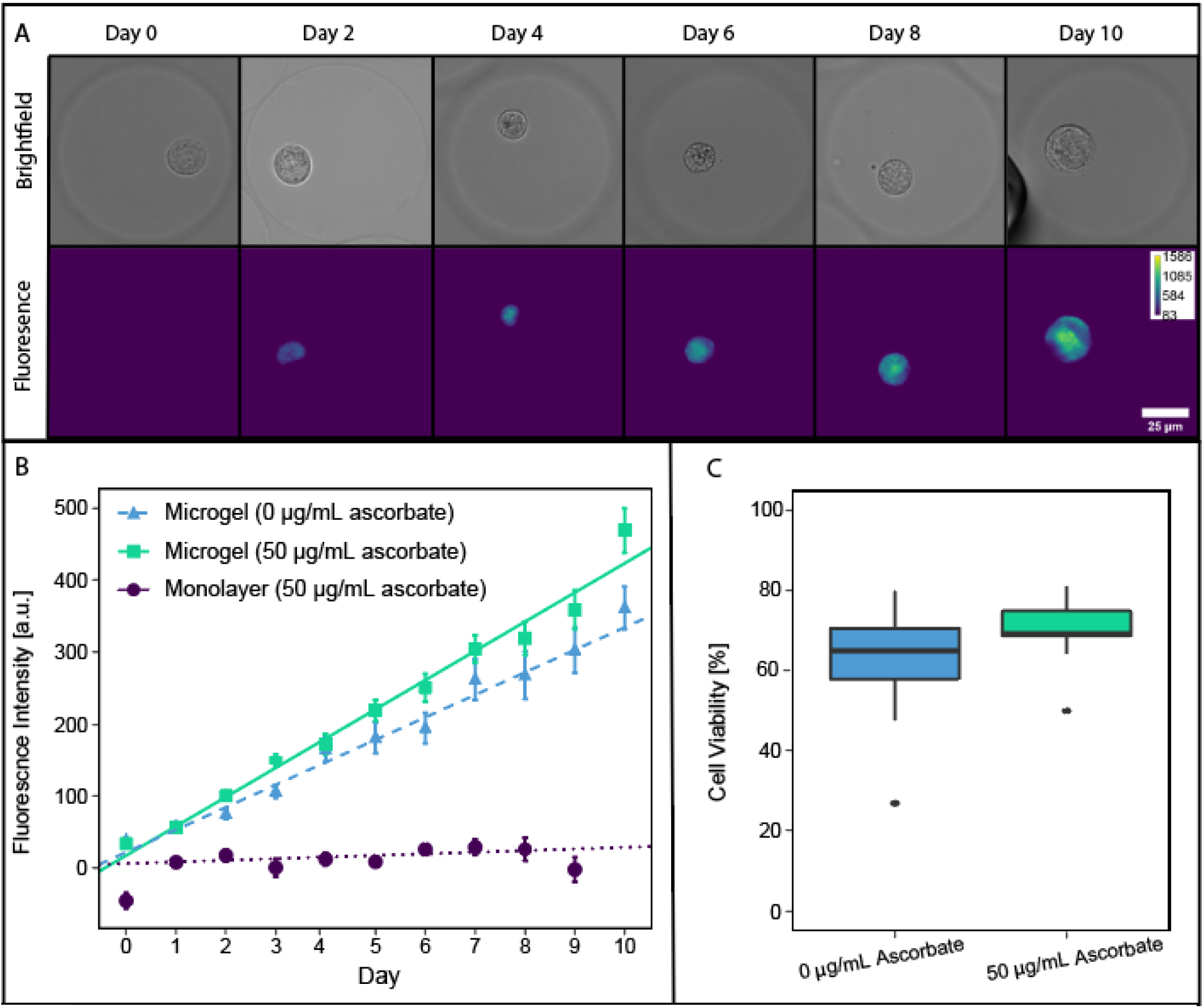
(A) Brightfield and fluorescence images (ex. 350/em. 470) of SW1353 cells encapsulated in alginate microgels and cultured with 50 μg/mL ascorbate. (B) Measured mean fluorescence intensities of imaged SW 1353 cells plotted with a weighted multiple linear regression. SW1353 cells encapsulated in microgels cultured without ascorbate (▴, -·-), encapsulated in microgels cultured with ascorbate (▪,---), monolayers cultured with ascorbate (●,···). Scale bar is 25 μm. (C) 10 day cell viability for SW1353 cells encapsulated in alginate microgels. Cell viability between cells cultured with or without ascorbate in microgels do not vary (p-value = 0.0957).

### Chondrocyte Viability in Alginate Microgels

After 10 days of culture in alginate microgels, SW1353 cell viability was evaluated by Trypan Blue exclusion. When 50 μg/mL ascorbate was present in the media, the cell viability was 70.3 ± 4.3%. Without ascorbate, the cell viability was 62.4 ± 7.7% (Fig. 3C). Using a Welch Two Sample t-test, no evidence was found to suggest a statistical difference in cell viability (p-value = 0.0957) indicating that 50 μg/mL of ascorbate likely does not affect SW1353 viability in alginate microgels.

### Elastic Moduli Characterization of Hydrogel Constructs

To quantify the mechanical properties of these hydrogels with and without embedded microgels, the storage and loss moduli of the 1.5% w/w alginate, the 4.5% w/w agarose, and the 4.5% w/w agarose with 20% v/v 1.5 % w/w alginate microgels were measured on a parallel plate rheometer (SI Table 1). The 1.5 % w/w alginate had storage and elastic moduli two orders of magnitude lower than the agarose (SI Fig. 3). There was no significant difference between the stiffness of the agarose gels with and without embedded alginate microgels.

### Metabolomic Profiling

To examine the effects of microgel encapsulation on chondrocyte mechanotransduction, SW1353 cells encapsulated in alginate microgels were suspended in a 4.5 % w/w agarose solution, poured into a cylindrical mold, and gelled (Fig. 1 B). 15 minutes of dynamic compression at 1.1 Hz was applied using a custom-built bioreactor (Fig. 1 C) [8]. Sample groups included both cells encapsulated in microgels before agarose encapsulation and control cells encapsulated only in agarose after monolayer culture. Using principle components analysis and volcano plots, distinct profiles between uncompressed monolayer and microgels were found using HPLC-MS (Figs. 4-5). Both hierarchical clustering and principal components analysis found substantial differences between metabolomic profiles from microgels and monolayer chondrocytes independent of compression (Fig. 4). There were substantial differences between metabolomic profiles of uncompressed microgel and uncompressed monolayer samples with 174 metabolites upregulated in microgels and 589 upregulated in monolayer (Fig. 5 A).Furthermore, there were substantial differences between the profiles of compressed microgels and compressed monolayer cells upon pairwise comparison (Fig. 5 B). Metabolomic profiles differed between compressed microgels and compressed monolayers with 75 metabolites upregulated in microgels and 399 metabolites upregulated in monolayer samples (Fig. 5 B). Clustering of median metabolite intensities found similarity between uncompressed microgels cultured with ascorbate and compressed microgels without ascorbate, indicating that compression can mimic the effects of ascorbate on chondrocyte metabolomic profiles (SI Figs. 4-8). Within both the monolayer and microgel groups, compression induced a robust metabolomic response for SW1353 chondrocytes (Fig. 5 C-D). Furthermore, compressed samples showed decreased variability in metabolomic profiles, as previously observed [8, 22].

**Figure 4:**
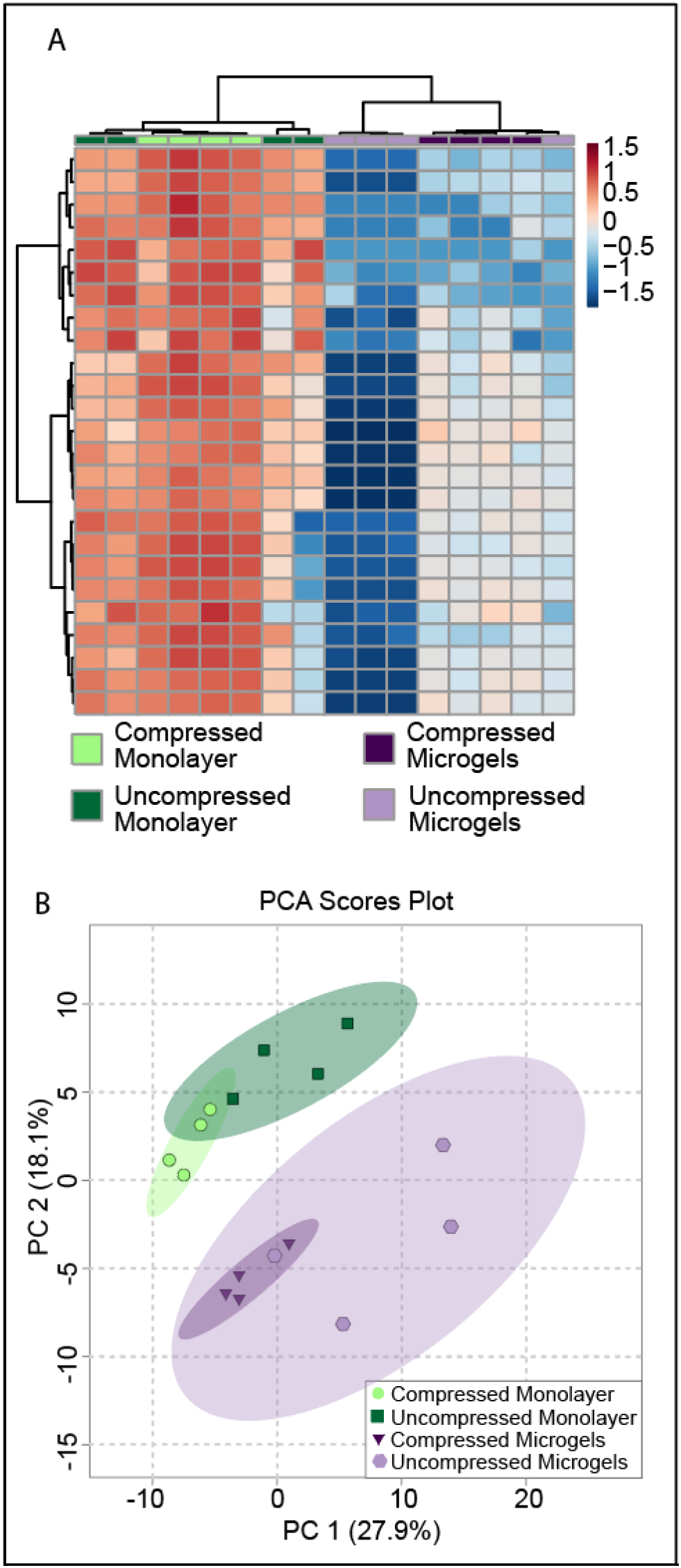
(A) Heat map of top 25 metabolites from SW1353 cells. (B) 2-D scores plot of PCA analysis for metabolites from SW1353 cells cultured with complete media supplemented with 50μg/ml L-sodium ascorbate.

**Figure 5:**
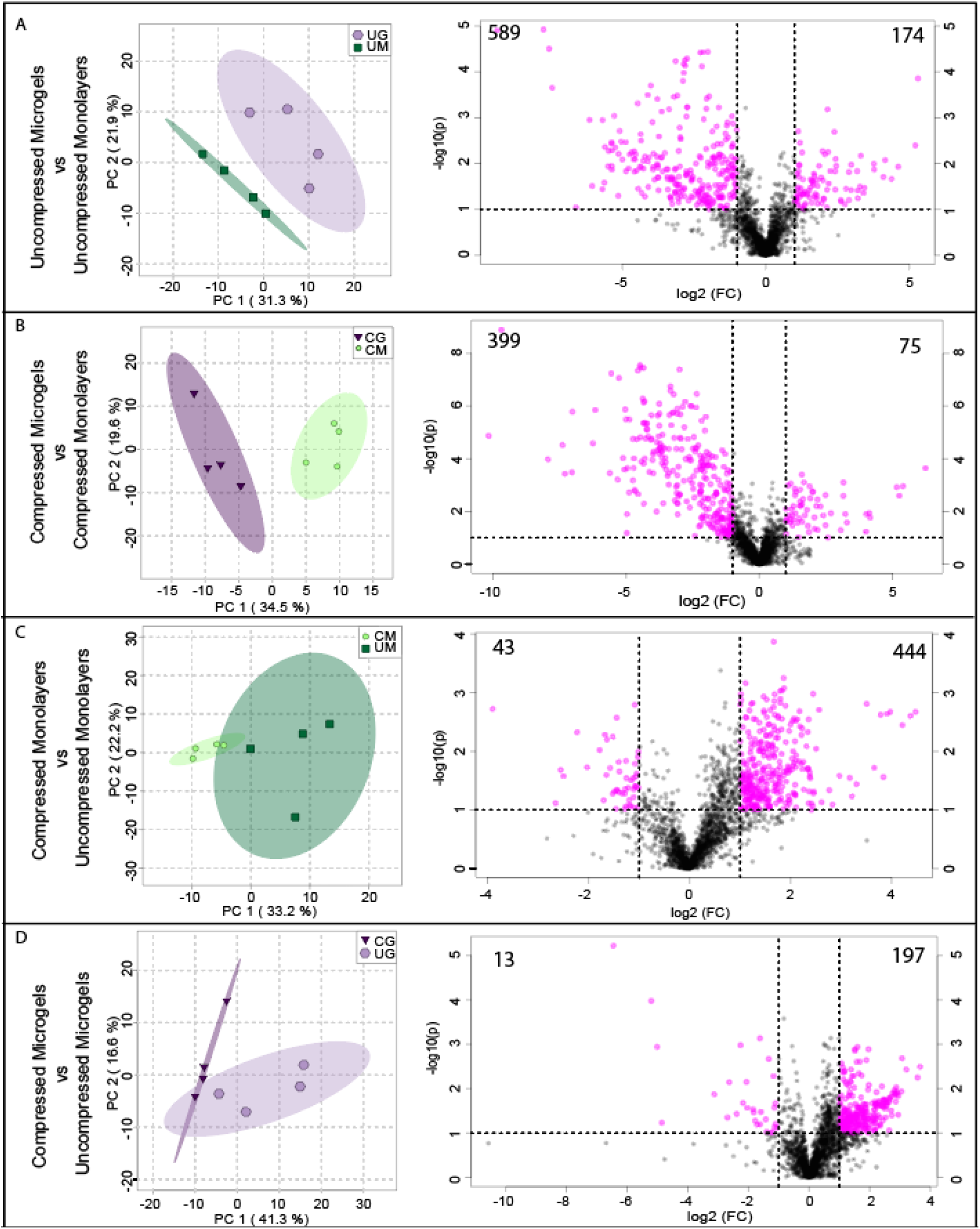
2-D score plots for PCA analysis (left) and volcano plots (right) for metabolites from SW1353 cells. (A) Uncompressed Monolayers (UM) vs. Uncompressed Microgels (UG). (B) Compressed Microgels (CG) vs. Compressed Monolayers (CM). (C) Uncompressed Monolayers vs. Compressed Monolayers. (D) Compressed Microgels vs. Uncompressed Microgels. All comparisons are made for cells cultured in complete media supplemented with 50 μg/ml L-sodium ascorbate. The numbers in the corners of the volcano plots denote the number of up and down regulated metabolites.

Dynamic compression in microgels induced upregulation of 197 metabolites and downregulation of 13 metabolites when compared with uncompressed controls (Fig. 5 D). Finally, microgels had distinct compression-induced responses from compressed monolayer controls. Pathway enrichment analysis of statistically-significant metabolites between groups did not find any pathways, potentially indicating broad changes in metabolite expression patterns between monolayer and microgel conditions. These data indicate that compression of alginate-encapsulated microgels results in a distinct metabolomic profile from compression of monolayer-expanded chondrocytes. Preculture in alginate microgels allows development of a type VI-rich pericellular matrix, similar to the *in vivo* structure that improves physiological *in vitro* modeling of chondrocyte mechanotransduction.

## Conclusions

Here chondrocytes were encapsulated in alginate microgels and imaged daily and evaluated for collagen formation. The data indicate that collagen VI production is greatly increased in microgels compared to monolayer cultures. Primary human chondrocytes produced more collagen VI than SW1353 cells when cultured in microgels or monolayers as observed using confocal microscopy. Thus, these results indicate that chondrocytes are more likely to form a PCM comprised of collagen VI in alginate microgels than in monolayers. Additionally, chondrocytes were encapsulated in physiologically stiff agarose gels and subjected to physiological dynamic compression. These cells cultured in microgels had substantially different metabolomic profiles compared to cells cultured in monolayers. Compression induced a robust metabolic response in both monolayer controls and microgel-encapsulated chondrocytes. These results provide the foundation for future studies on chondrocyte mechanotransduction. By encapsulating single chondrocytes in alginate microgels, PCM formation is more likely to be induced and will alter how chondrocytes respond to physiological dynamic compression.

In summary, we show that microgels created using drop-based microfluidics can provide a robust 3D culture environment for individual chondrons *in vitro*. By encapsulating the cells individually in small volumes of alginate, microgels allow for the manipulation of single cells in their native microenvironments. This novel method eliminates the use of potentially harmful digestive processes that are typically applied to isolate single cells in traditional bulk alginate culture. This is the first instance of chondrons encapsulated in alginate microgels establishing methods for future work on primary human chondrocyte mechanotransduction and chondrocyte tissue engineering.

## Acknowledgments

This work was supported by the National Institutes of Health [R01AR073964 to RKJ], National Science Foundation [CAREER 1753352 to CBC, 1554708 to RKJ]. The funding sources had no role in the design or execution of the study.

## Data availability

The authors declare that all data presented in this manuscript and all other relevant data supporting the findings of this study are available from the corresponding author upon reasonable request.

## Supplemental Figures and Tables

**SI Figure 1:**
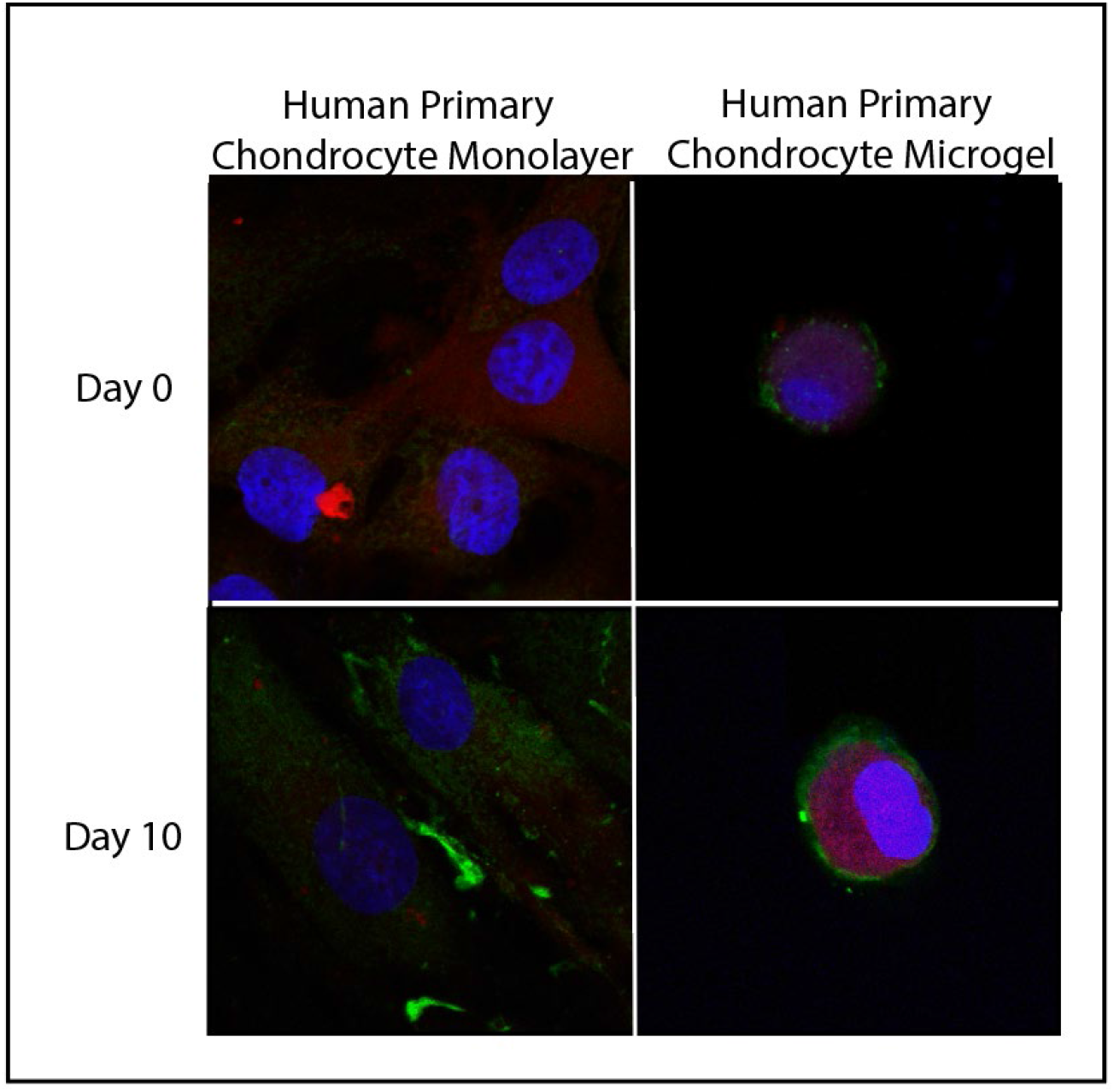
Primary human chondrocyte cells fixed with 4% v/v paraformaldehyde in PBS. Cells are stained for Collagen VI (green), mitochondria (red), and the nuclei (blue). Cells were imaged on days 0 and 10 on monolayers and encapsulated in 1.5% w/w alginate microgels. Cells pictured were cultured in media without ascorbate.

**SI Figure 2:**
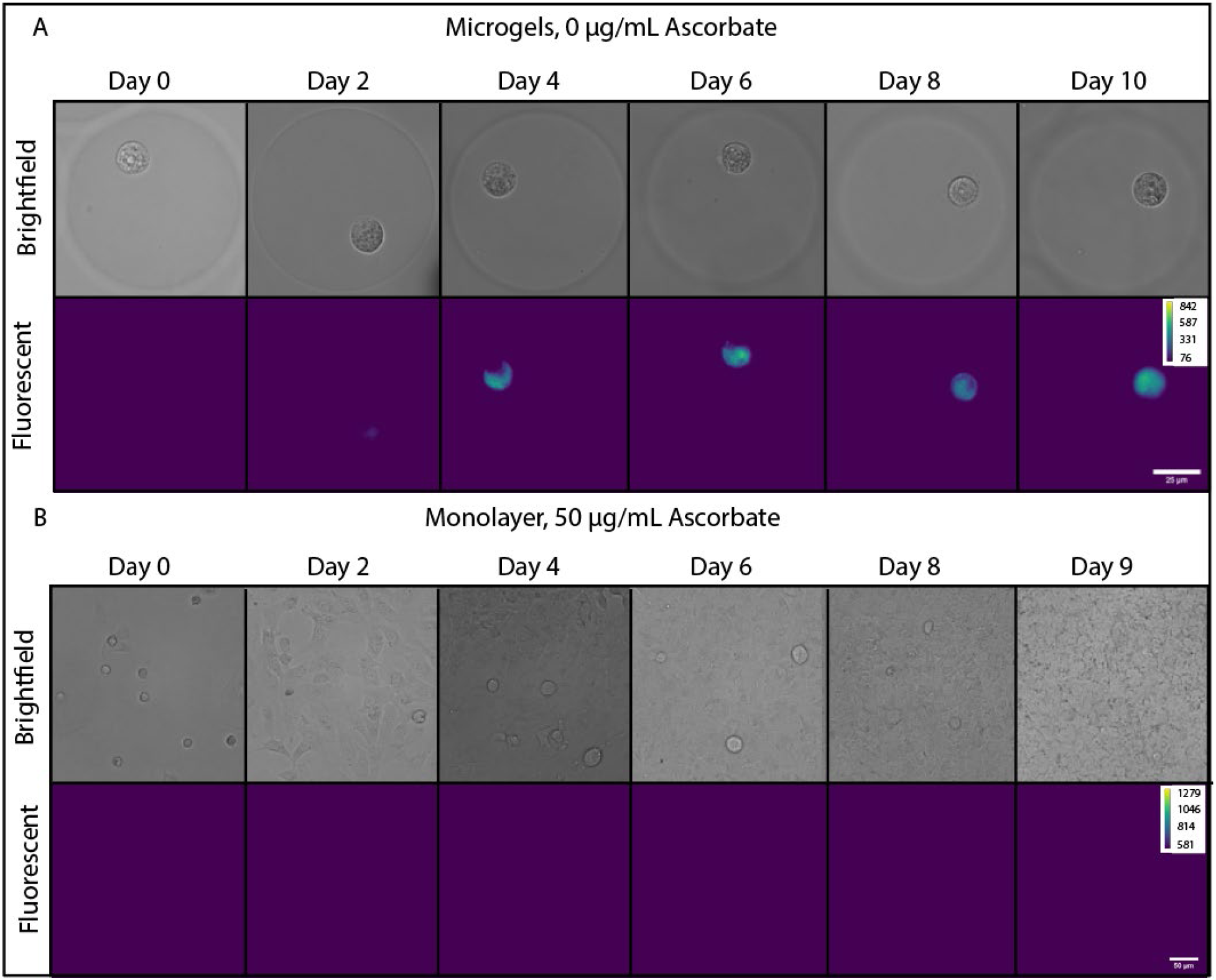
Microgels without and with ascorbate (a) Brightfield and fluorescence images of SW1353 cells encapsulated in alginate microgels and cultured without ascorbate. (b) Brightfield and fluorescence images of SW1353 cells grown on monolayers with 50 μg/mL ascorbate.

**SI 3:**
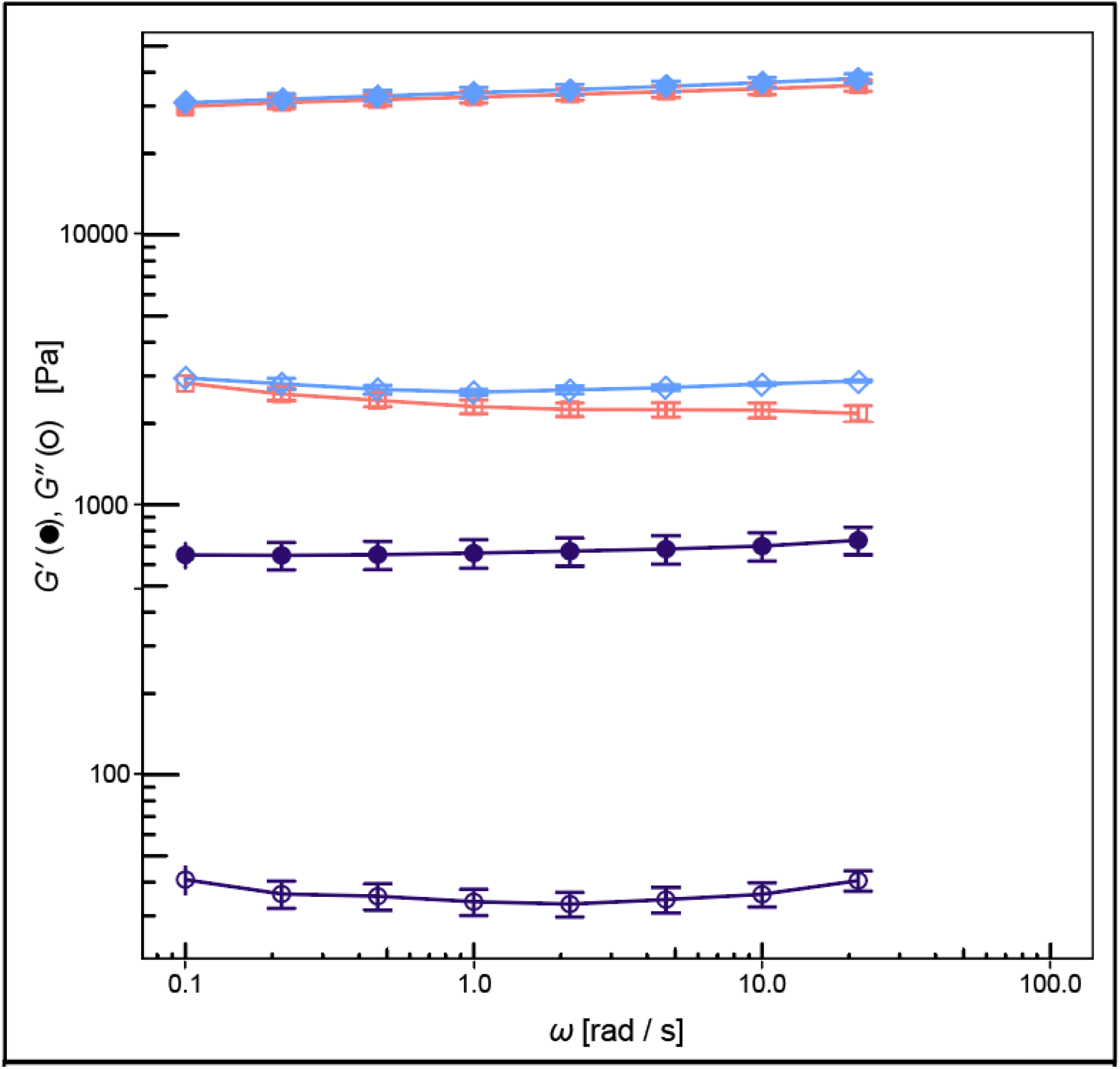
Storage (closed symbols) and loss (open symbols) moduli versus angular frequency, *T* = 25 °C, for 4.5% w/w agarose (squares), 1.5% w/w alginate (circles), and 4.5% w/w agarose with 20% v/v 1.5% w/w alginate microgels (diamonds).

**SI Figure 4:**
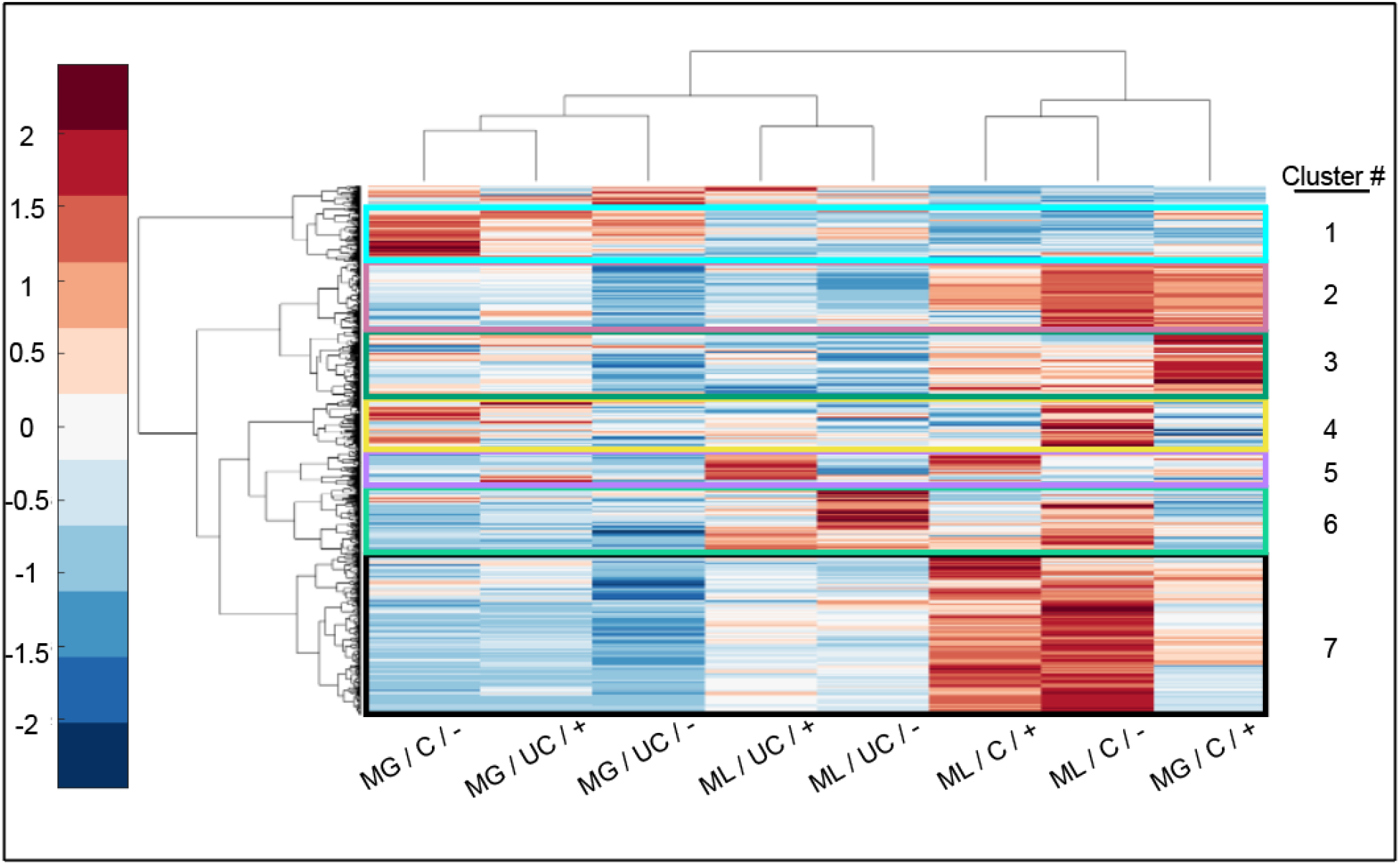
Median metabolites across all experimental groups. Metabolite clusters (colored boxes) were subjected to pathway enrichment analysis, but no significant pathways were found after false discovery rate correction. ML (Monolayer), MG (Microgel), C (Compressed), UC (Uncompressed), + (complete media supplemented with 50 μg/mL L-sodium ascorbate), - (complete media lacking L-sodium ascorbate).

**SI Figure 5:**
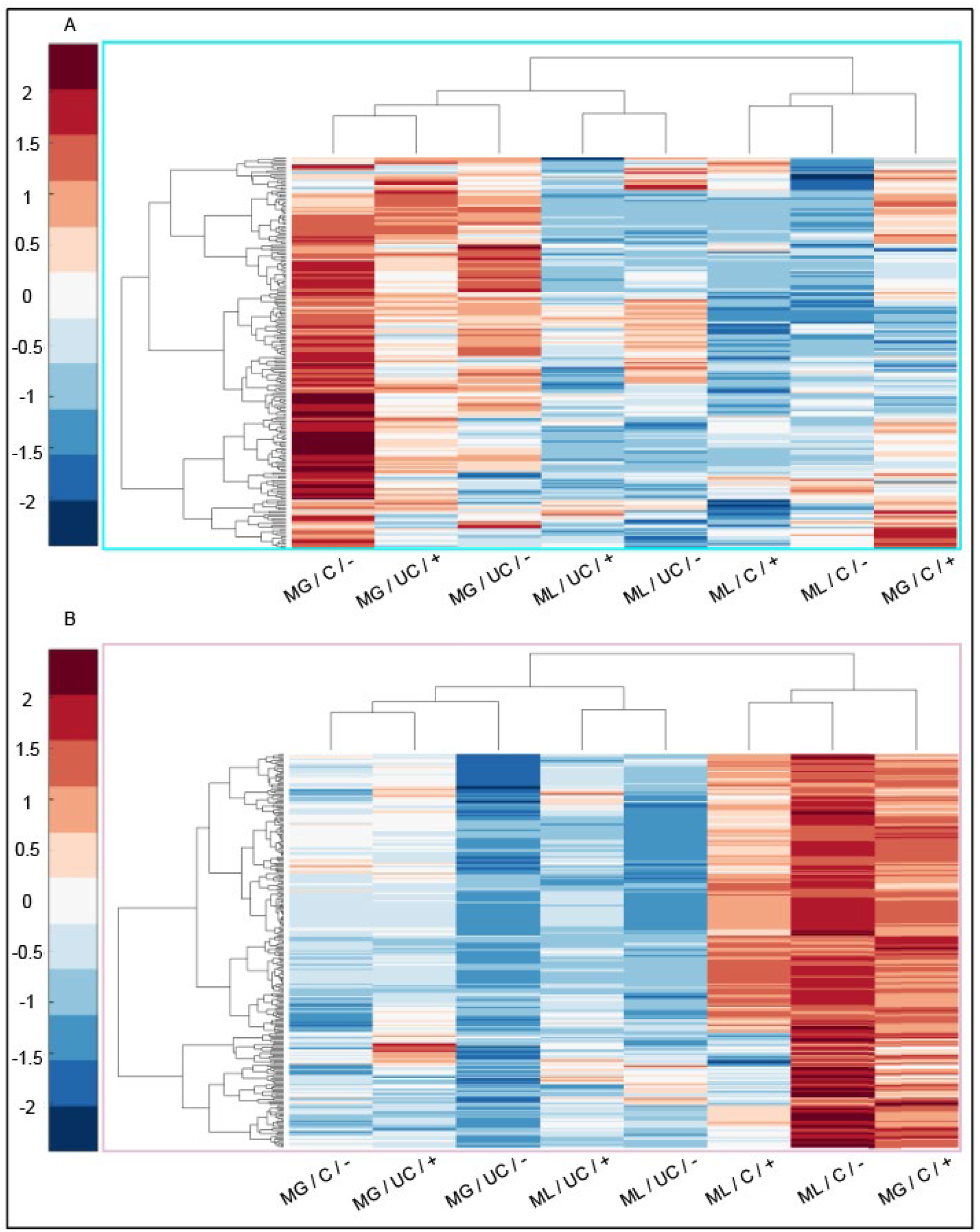
Median metabolites across all experimental group. (A) Metabolite cluster 1 and (B) cluster 2 from SI figure 4 were further subjected to pathway enrichment analysis, but no significant pathways were found after false discovery rate correction. ML (Monolayer), MG (Microgel), C (Compressed), UC (Uncompressed), + (complete media supplemented with 50 μg/mL L-sodium ascorbate), - (complete media lacking L-sodium ascorbate).

**SI Figure 5:**
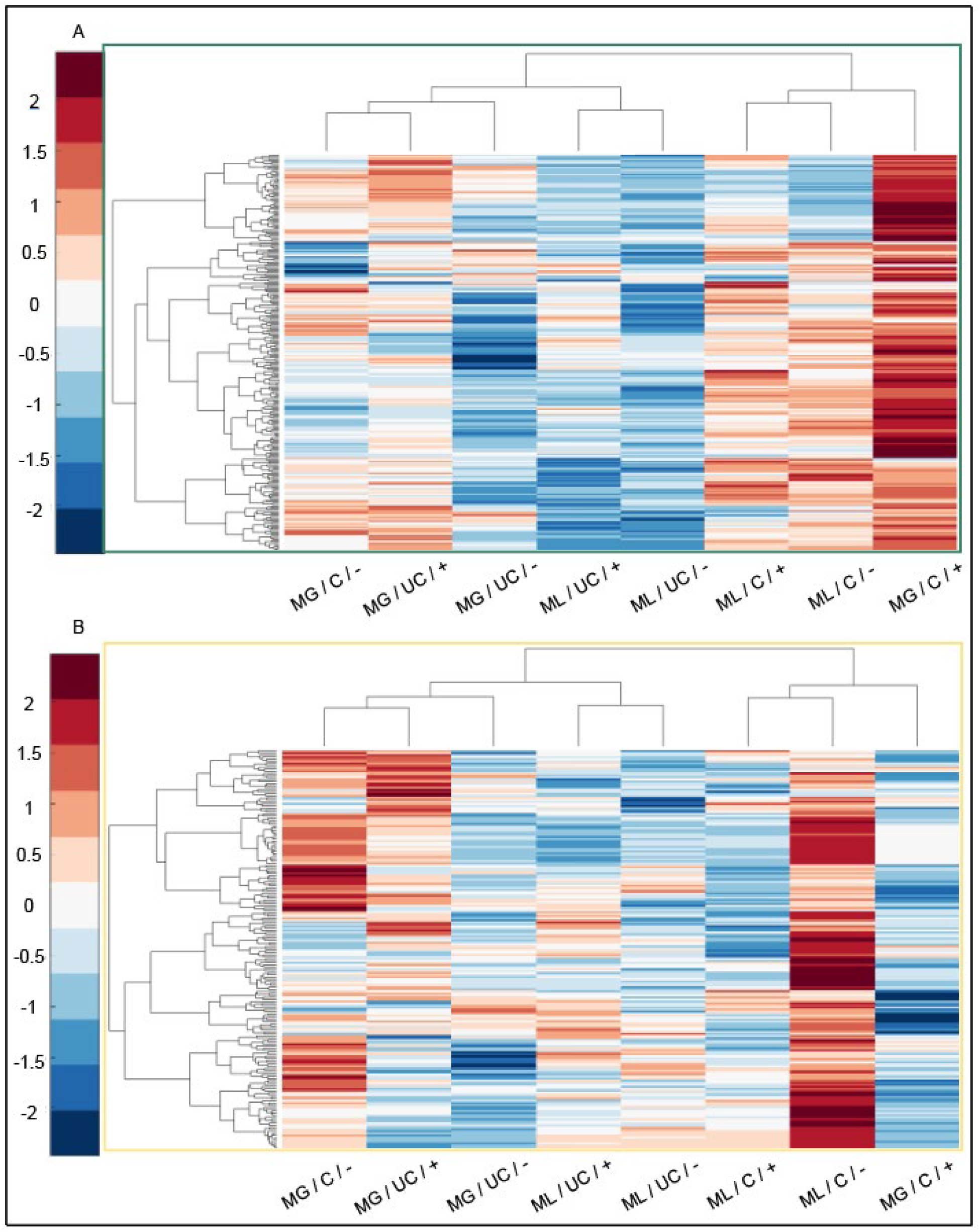
Median metabolites across all experimental group. (A) Metabolite cluster 2 and (B) cluster 3 from SI Figure 4 were subjected to pathway enrichment analysis, but no significant pathways were found after false discovery rate correction. ML (Monolayer), MG (Microgel), C (Compressed), UC (Uncompressed), + (complete media supplemented with 50 μg/mL L-sodium ascorbate), - (complete media lacking L-sodium ascorbate).

**SI Figure 6:**
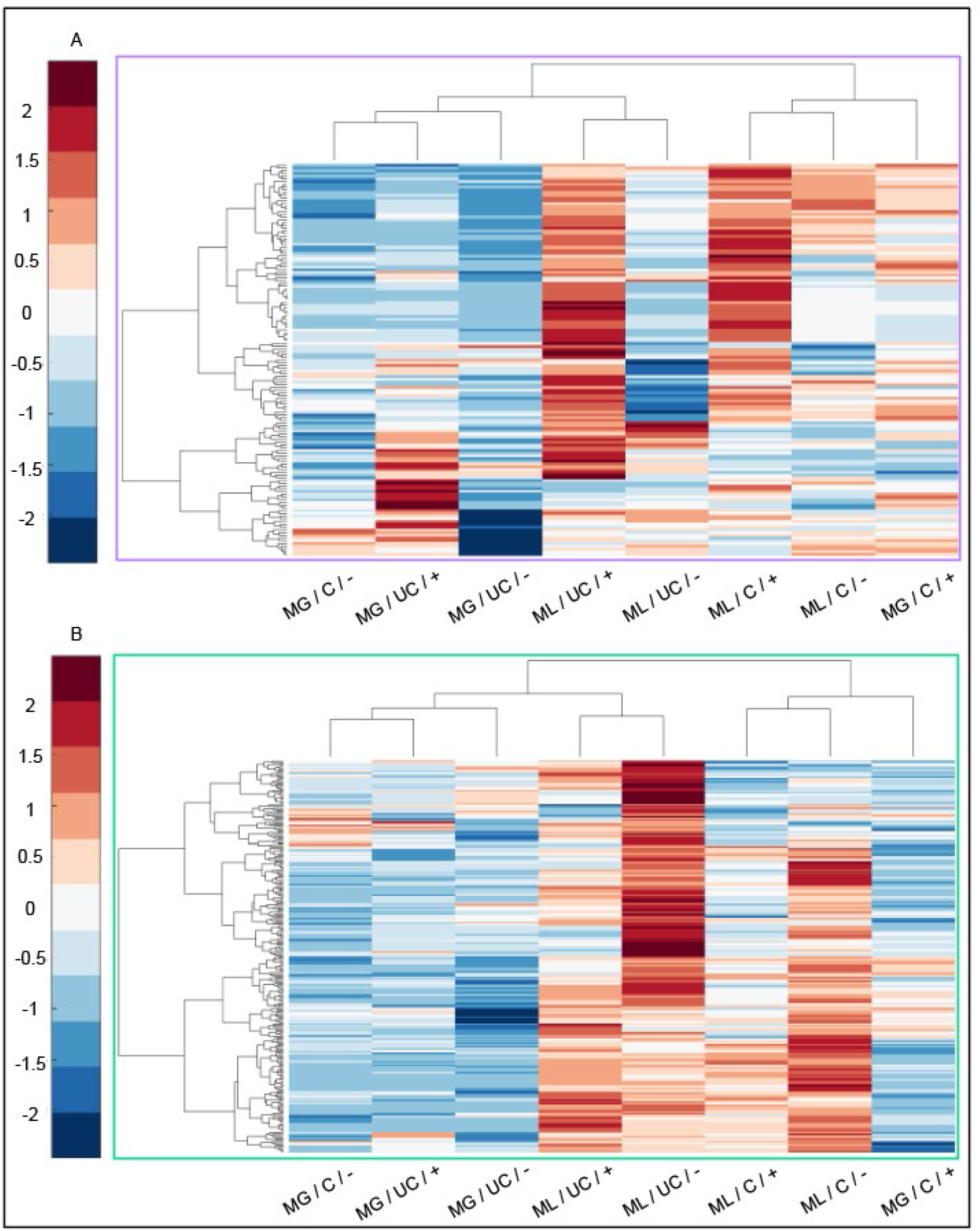
Median metabolites across all experimental group. (A) Metabolite cluster 5 and (B) cluster 6 from SI Figure 4 were subjected to pathway enrichment analysis, but no significant pathways were found after false discovery rate correction. ML (Monolayer), MG (Microgel), C (Compressed), UC (Uncompressed), + (complete media supplemented with 50 μg/mL L-sodium ascorbate), - (complete media lacking L-sodium ascorbate).

**SI Figure 7:**
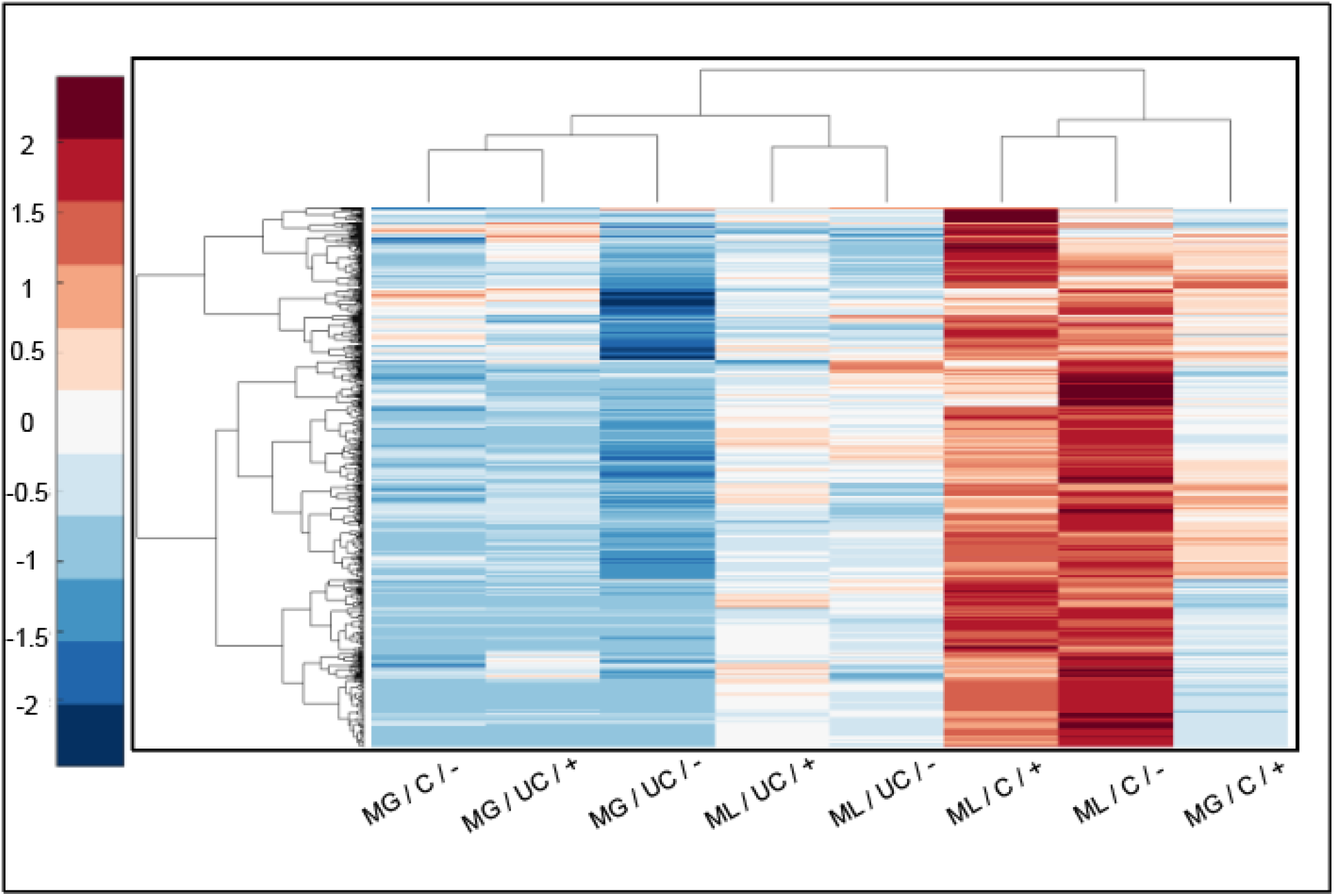
Median metabolites across all experimental group. Metabolite cluster 7 from SI Figure 4 was subjected to pathway enrichment analysis, but no significant pathways were found after false discovery rate correction. ML (Monolayer), MG (Microgel), C (Compressed), UC (Uncompressed), + (complete media supplemented with 50 μg/mL L-sodium ascorbate), - (complete media lacking L-sodium ascorbate).

**SI T1:** Elastic (G’) and Storage (G’’) moduli of 4.5% w/w agarose, 1.5% w/w alginate, and 4.5 % w/w agarose with 20% v/v 1.5% w/w alginate microgels for varying angular frequencies. Rheology was performed on a TA Instruments AR-G2 rheometer with a parallel plate geometry.

| Hydrogel | $\omega$ [rad/s] | $G'$ [Pa] | | $G''$ [Pa] | |
| --- | --- | --- | --- | --- | --- |
| 4.5% w/w Agarose | 0.100 | 3.0E+04 | ± 2.9E+03 | 2.8E+03 | ± 2.4E+02 |
|  | 0.215 | 3.1E+04 | ± 2.7E+03 | 2.6E+03 | ± 2.4E+02 |
|  | 0.464 | 3.2E+04 | ± 2.8E+03 | 2.4E+03 | ± 2.2E+02 |
|  | 1.000 | 3.2E+04 | ± 2.8E+03 | 2.3E+03 | ± 2.3E+02 |
|  | 2.154 | 3.3E+04 | ± 2.8E+03 | 2.3E+03 | ± 2.2E+02 |
|  | 4.642 | 3.4E+04 | ± 2.9E+03 | 2.2E+03 | ± 2.4E+02 |
|  | 10.000 | 3.5E+04 | ± 3.0E+03 | 2.2E+03 | ± 2.4E+02 |
|  | 21.544 | 3.6E+04 | ± 3.0E+03 | 2.2E+03 | ± 2.6E+02 |
| 1.5 % w/w Alginate | 0.100 | 6.5E+02 | ± 1.8E+02 | 4.1E+01 | ± 1.2E+01 |
|  | 0.215 | 6.5E+02 | ± 1.9E+02 | 3.6E+01 | ± 1.0E+01 |
|  | 0.464 | 6.5E+02 | ± 1.9E+02 | 3.5E+01 | ± 9.8E+00 |
|  | 1.000 | 6.6E+02 | ± 2.0E+02 | 3.4E+01 | ± 9.3E+00 |
|  | 2.154 | 6.7E+02 | ± 2.0E+02 | 3.3E+01 | ± 8.5E+00 |
|  | 4.642 | 6.9E+02 | ± 2.0E+02 | 3.4E+01 | ± 9.2E+00 |
|  | 10.000 | 7.0E+02 | ± 2.1E+02 | 3.6E+01 | ± 9.1E+00 |
|  | 21.544 | 7.4E+02 | ± 2.1E+02 | 4.1E+01 | ± 8.6E+00 |
| Agarose + 20% v/v Alginate Microgels | 0.100 | 3.1E+04 | ± 2.7E+03 | 2.9E+03 | ± 1.6E+02 |
|  | 0.215 | 3.2E+04 | ± 2.8E+03 | 2.8E+03 | ± 2.3E+02 |
|  | 0.464 | 3.3E+04 | ± 2.7E+03 | 2.7E+03 | ± 1.6E+02 |
|  | 1.000 | 3.4E+04 | ± 2.7E+03 | 2.6E+03 | ± 1.0E+02 |
|  | 2.154 | 3.4E+04 | ± 2.7E+03 | 2.7E+03 | ± 1.5E+02 |
|  | 4.642 | 3.5E+04 | ± 2.7E+03 | 2.7E+03 | ± 1.1E+02 |
|  | 10.000 | 3.7E+04 | ± 2.7E+03 | 2.8E+03 | ± 5.9E+01 |
|  | 21.544 | 3.8E+04 | ± 2.7E+03 | 2.9E+03 | ± 4.2E+01 |

